## Supplementary Text for "Cloud gazing: demonstrating paths for unlocking the value of cloud genomics through cross-cohort analysis"

Methods

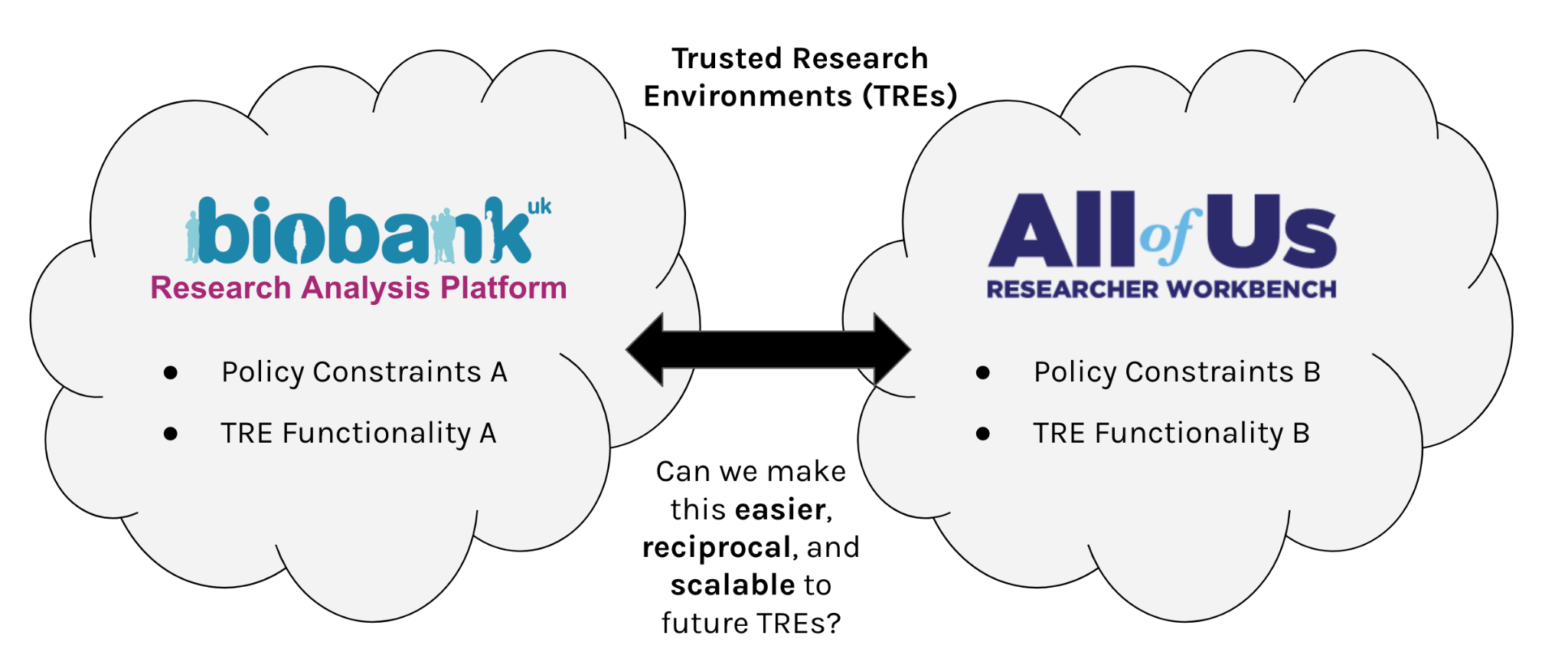

**Supplemental Fig 1.** Current state and important considerations to enable cross-cohort analysis of genomic data stored in separate Trusted Research Environments (TREs).

### **Data access**

#### **Policy details of data access**

The UK Biobank, a population-based cohort of approximately 500,000 participants recruited from 2006 to 2010, has existing genomic and longitudinal phenotypic data. Baseline assessments were conducted at 22 assessment centers across the United Kingdom, with sample collections including blood-derived DNA. UK Biobank individual-level data are available by request via application (<https://www.ukbiobank.ac.uk>). All UK Biobank participants gave written, informed consent per the UK Biobank primary protocol. Secondary use of this data was approved by the Massachusetts General Hospital Institutional Review Board (protocol 2021P002228) and was facilitated through UK Biobank application 7089.

Authorization for access to participant-level data in *All of Us* is user based, rather than project based. Authorized users receive a “data passport.” The data passport is required for gaining data access and for creating workspaces to carry out research projects using *All of Us* genomic and phenotypic data. Data passports are currently authorized via a 6-step process that includes affiliation with an institution which has signed a Data Use and Registration Agreement, account creation, identity verification using login.gov, completion of ethics training, and attestation to the *All of Us* Data User Code of Conduct. Additionally, approval to use the dataset for program operational demonstration projects was obtained from the *All of Us* Institutional Review Board. The *All of Us* Data and Statistics Dissemination Policy disallows disclosure of participant counts under 20 and allele counts under 40 to protect participant privacy (<https://www.researchallofus.org/data-tools/data-access/>) but this study applied for and received an exception to the policy.

#### **Technical details of data access**

For the UK Biobank analyses, a DNAnexus project for Broad UK Biobank application 7089 project created on the UK Biobank Research Analysis Platform.

For the *All of Us* analyses, a workspace was created on the *All of Us* Researcher Workbench.

For the pooled analyses, a workspace was also created on the *All of Us* Researcher Workbench. Broad UK Biobank application 7089 already had data staged in [Terra](https://app.terra.bio/) on Google Cloud for use in other research projects. The Broad data custodian added our @researchallofus.org accounts to the access control list for read-only access to the UK Biobank data so that the data became visible to the workspace within the *All of Us* Researcher Workbench (**Supplemental Fig. 2**).

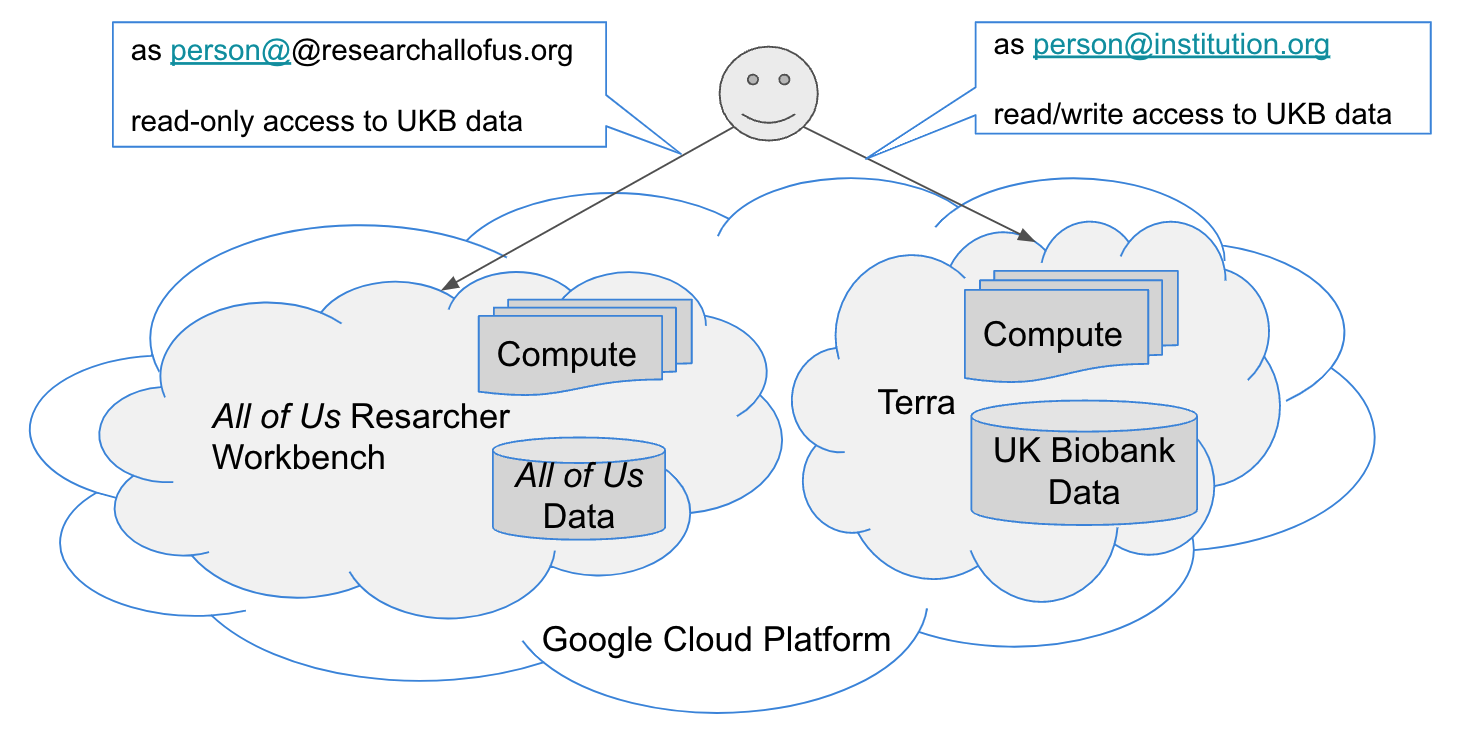

***Supplemental Fig. 2****: Data locations and access control configuration for the pooled analyses.*

### **Code availability**

The code for all analyses can be found in <https://github.com/all-of-us/ukb-cross-analysis-demo-project> and was compatible with UK Biobank Research Analysis Platform and *All of Us* Researcher Workbench available data and technical capabilities as of the Spring of 2022.

### **Genomic data preparation and quality control**

UK Biobank’s 200k exome release ^1^ was chosen for use in this project because it is the most recent release of genomic data permitted by UK Biobank policy rules to be analyzed outside of the UK Biobank Research Analysis Platform. UK Biobank exome data includes ~10M exonic variants, with an average coverage of 20X in 95.6% of the sites. No variant- or sample-level filters were pre-applied to the pVCF or PLINK files. For the meta-analyses, the PLINK format files were filtered to variants within the exonic capture regions (<https://biobank.ndph.ox.ac.uk/ukb/refer.cgi?id=3803>) and variants with an alternate allele frequency of 6 or more. For the pooled analyses, the pVCF files were ingested into Hail (Hail Team. Hail 0.2.13-81ab564db2b4. <https://github.com/hail-is/hail/releases/tag/0.2.13>) and emitted as a matrix table similarly filtered to exonic capture regions. The data was then further filtered via plink to include only variants with an alternate allele frequency of 6 or more.

The *All of Us* whole genome sequence alpha 3 release is the very first release of genomic data available for the *All of Us* cohort. The *All of Us* policy rules require that this data may only be used within the *All of Us* Researcher Workbench. Whole genome sequence (WGS) data was generated on consented *All of Us* participants via a College of American Pathologists (CAP) / Clinical Laboratory Improvement Amendments (CLIA) validated pipeline. A count-called SNP/Indel joint call set was generated from WGS data according to GATK Best Practices. WGS data were provided in pVCF and Hail matrix table formats. For the meta-analyses, the matrix table was filtered to variants within the UK Biobank exonic capture regions and variants with an alternate allele frequency of 6 or more. For the pooled analyses, the matrix table was similarly filtered to exonic capture regions. The data was then further filtered via plink to include only variants with an alternate allele frequency of 6 or more. We were not able to use the provided VCF filter flags for filtering since each cohort used different flagging criteria (**Supplemental Fig. 3**).

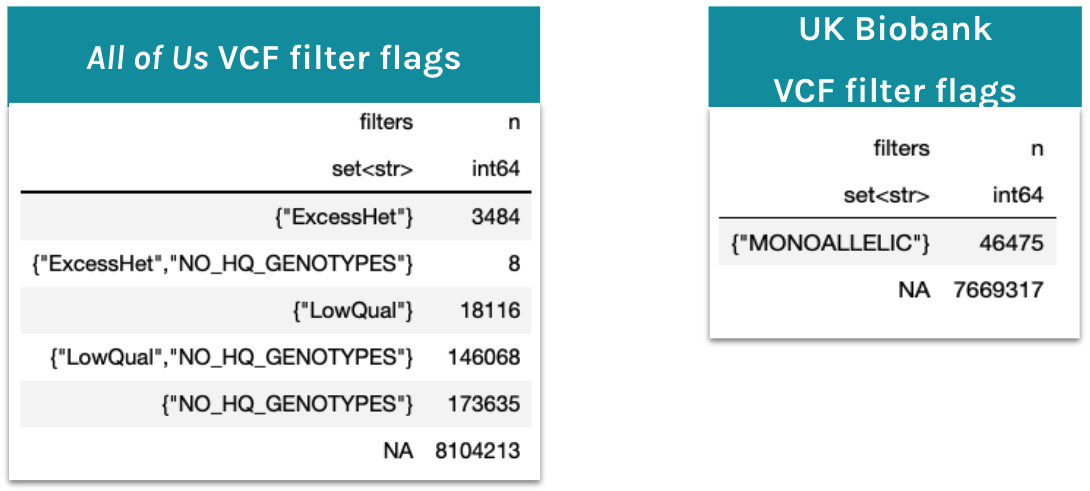

***Supplemental Fig. 3****: The UK Biobank and All of Us VCF data used* ***different*** *soft thresholds in the VCF filter field, therefore we were not able to use these precomputed results in our filtering.*

For the pooled analysis, biallelic variants were merged if these values were identical: [chrom, pos, ref, alt] (**Supplemental Fig. 4a and 4b**). More specifically, the variants in the prepared UK Biobank and *All of Us* matrix tables were split into biallelic variants using Hail method [split_multi_hts](https://hail.is/docs/0.2/methods/genetics.html#hail.methods.split_multi_hts) and then an inner join of the variants was performed via Hail method [union_cols](https://hail.is/docs/0.2/hail.MatrixTable.html#hail.MatrixTable.union_cols). For full details, please see [03_merge_variants.ipynb](https://github.com/all-of-us/ukb-cross-analysis-demo-project/blob/main/aou_workbench_pooled_analyses/03_merge_variants.ipynb).

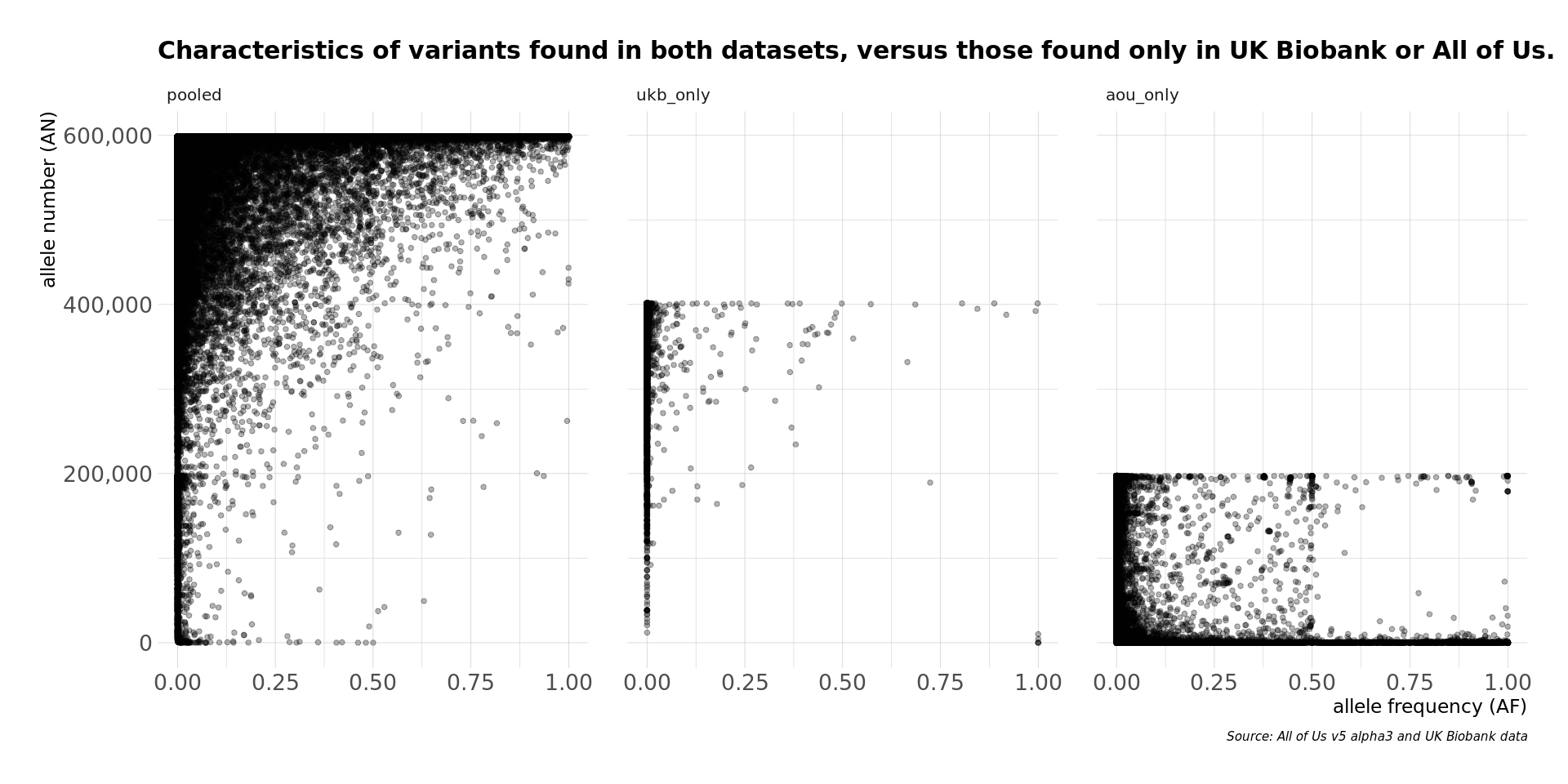

***Supplemental Fig. 4a****: The maximum value for allele number on the y-axis is determined by cohort size. The pooled exonic variants consist of common and rare variants. Most exonic variants found in UK Biobank only were very rare. Most exonic variants found in All of Us only were either very rare or have a very low allele number. Note that variant QC and AC filtering has not yet occurred for the data shown in these plots. From the All of Us VCF filter field values for these variants, most of the common variants found in All of Us only were of low quality and would have eventually been filtered out during variant QC, if they had been included.*

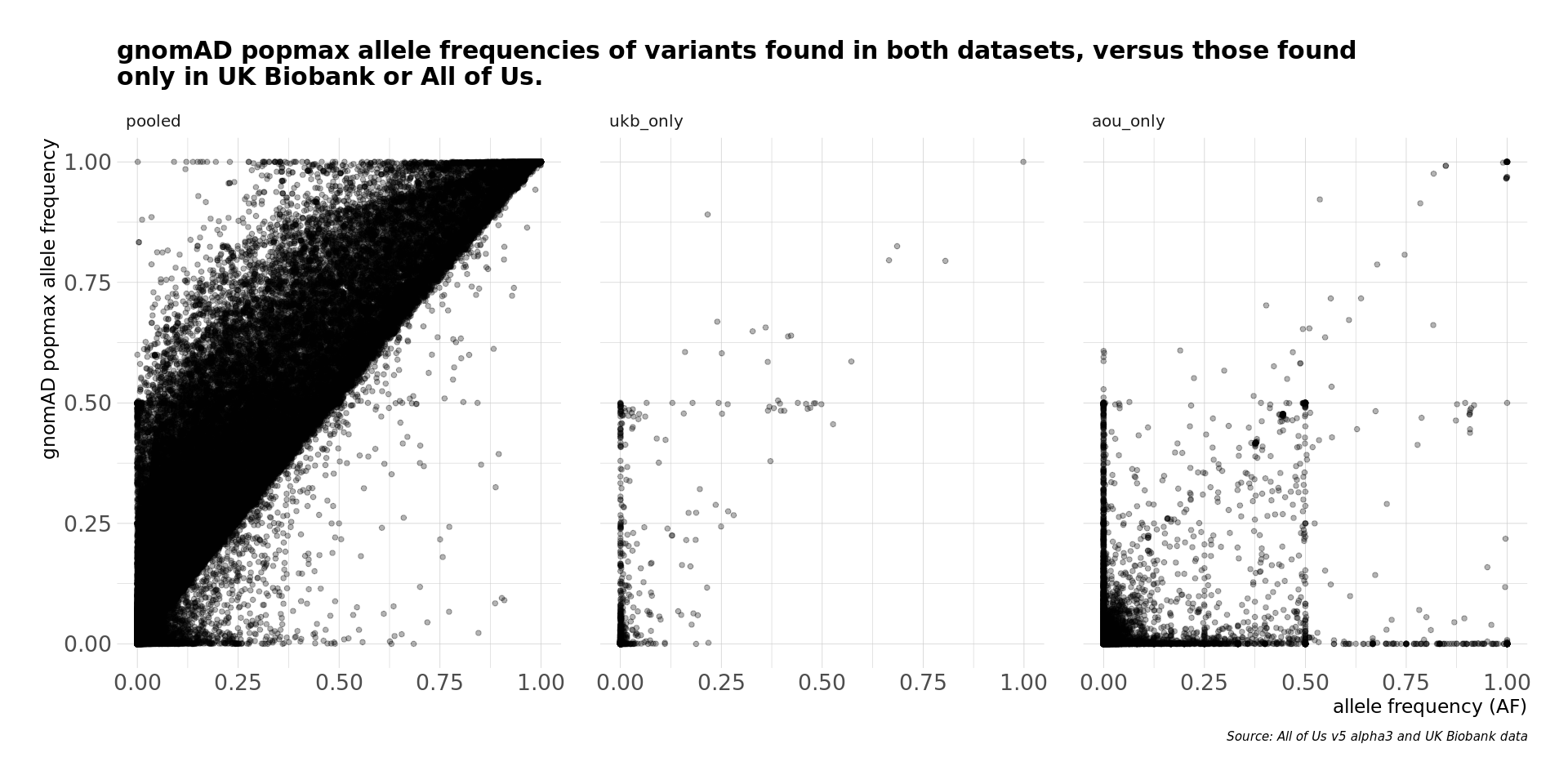

***Supplemental Fig. 4b****: The pooled exonic variants show a clear pattern with gnomAD population maximum allele frequencies. Less alignment along the diagonal is shown for UK Biobank only and All of Us only exonic variants. Note that variant QC and AC filtering has not yet occurred for the data shown in these plots. From the All of Us VCF filter field values for these variants, most of the common variants found in All of Us only were of low quality and would have eventually been filtered out during variant QC, if they had been included.*

Variants were removed from use in downstream analyses, such as principal component analysis and REGENIE, if Hardy-Weinberg equilibrium exact test p-value was below 1e-15 or missing call rates exceed 10%. Samples were removed if missing call rates exceeded 10%, but no samples in UK Biobank or *All of Us* exceeded this missingness threshold. We did not apply other variant QC criteria such as call quality thresholds because the determination of equivalent thresholds for use with DeepVariant+GLnexus variants versus DRAGEN variants is non-trivial to determine due to their differences in accuracy ^2^.

### **Phenotype preparation**

Blood lipids including low-density lipoprotein cholesterol (LDL-C), total cholesterol (TC), high-density lipoprotein cholesterol (HDL-C) and triglycerides (TG) were used as the primary phenotypes in this study. We curated and harmonized the lipid measurements and statin drug exposures for both UK Biobank and *All of Us* from the phenotype resources of these cohorts.

#### **Lipid levels**

For UK Biobank, study-specific blood serum lipids assays were performed systematically in its central laboratories^3^. The first instance of the lipid measurement was used which included LDL-C (data-field-ID: 30780), TC (data-field-ID: 30690), HDL-C (data-field-ID: 30760) and TG (data-field-ID: 30870). Most participants (N=190,982) in the 200k exome release had at least one non-null lipid value, and therefore were included in the analysis. The lipid measurements were converted from mmol/L to mg/dL by multiplying TG values by 88.57 and the other lipid measurements by 38.67 ^4^.

For *All of Us*, there were no study- specific blood serum lipids assays available at the time of this analysis, so we instead used lipids measurements from Electronic Health Records (EHR). Of the 98,622 WGS samples, 37,754 of the participants had at least one type of lipid measurement. The most recent measurement value for each of the four lipid types was used. Note that a person’s measurements for the four different lipid types, if available, may have occurred on different dates. In order to maximize the number of *All of Us* genomes we were able to include in this study, we collapsed several related OMOP measurement concepts and several OMOP unit concepts (**Supplemental Fig. 5**). This included use of measurements with no unit specified, when the data distribution for that measurement appeared empirically to be in mg/dL.

UNIT_NAMES **<-** c('milligram per deciliter', 'No matching concept', 'mg/dL')

*# HDL cholesterol*

HDL_MEASURE_NAMES **<-** c(

'Cholesterol in HDL [Mass/volume] in Serum or Plasma',

'Cholesterol in HDL [Mass/volume] in Serum or Plasma by Electrophoresis',

'Cholesterol in HDL [Mass/volume] in Serum or Plasma ultracentrifugate')

*# LDL-C cholesterol*

LDL-C_MEASURE_NAMES **<-** c(

'Cholesterol in LDL-C [Mass/volume] in Serum or Plasma by calculation',

'Cholesterol in LDL-C [Mass/volume] in Serum or Plasma',

'Cholesterol in LDL-C [Mass/volume] in Serum or Plasma ultracentrifugate',

'Cholesterol in LDL-C [Mass/volume] in Serum or Plasma by Direct assay',

'Cholesterol in LDL-C [Mass/volume] in Serum or Plasma by Electrophoresis')

*# Total cholesterol*

TC_MEASURE_NAMES **<-** c('Cholesterol [Mass/volume] in Serum or Plasma')

*# Triglycerides*

TG_MEASURE_NAMES **<-** c(

'Triglyceride [Mass/volume] in Serum or Plasma',

'Triglyceride [Mass/volume] in Blood',

'Triglyceride [Mass/volume] in Serum or Plasma --fasting',

'Triglyceride [Mass/volume] in Serum or Plasma by calculation')

***Supplemental Fig. 5****: Several OMOP measurement and unit concepts were collapsed into the final lipid phenotype.*

#### **Age at time of measurement**

For UK Biobank participants, their age at time of measurement was their age when they attended the assessment center (data-field-ID: 21003).

For *All of Us* participants, as noted in the prior section, a person’s measurements for the four different lipid types, if available, may have occurred on different dates. REGENIE can analyze multiple phenotypes concurrently, so a single value for the age covariate was desired. To achieve this single value for age, we used the maximum age at time of measurement determined from the person’s four, or fewer, different lipid type measurements. If any particular lipid type measurement was taken at an age more than 5 years earlier than their maximum age, it was discarded.

#### **Statin drug classification**

For UK Biobank, the generic and trade names of drugs were expressed in the controlled vocabulary Read v2 and Read v3.

For *All of Us*, the generic and trade names of drugs were expressed in controlled vocabulary RxNORM ^5^.

Note that since UK Biobank and *All of Us* used different controlled vocabularies for drugs, we were not able to utilize an identical set of statin drugs (**Supplemental Fig. 6**).

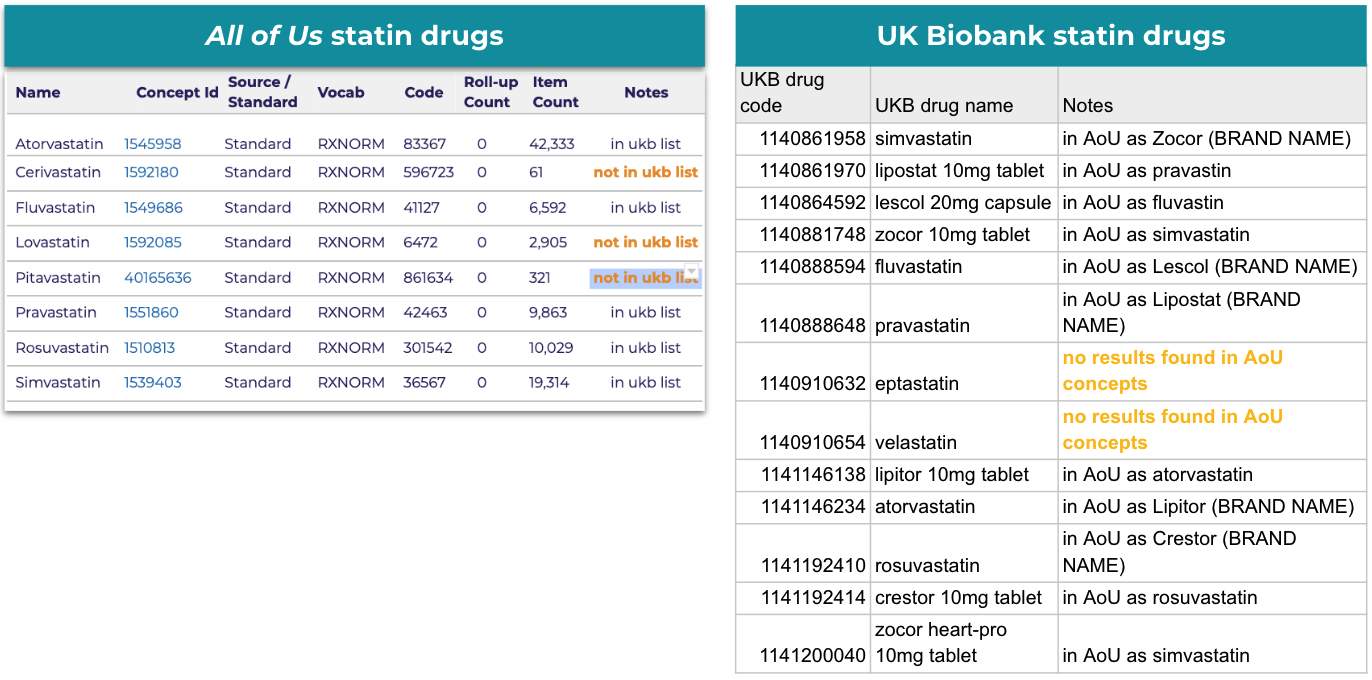

***Supplemental Fig. 6****: Left: the statin drug list used for All of Us statin use determination. Right: the statin drug list used for UK Biobank statin drug use determination.*

#### **Statin drug use determination**

For UK Biobank participants, a current medication list was provided in a verbal interview during their visit to the study center (data-field-ID 20003).

For *All of Us* participants, their visit to the study center had no specific relationship to drug exposures at the time of measurement, because the lipids measurements were from their historical EHR, not a study-specific blood serum assay. Therefore, we employed a heuristic method to determine whether a statin drug exposure was likely in effect when their lipids measurement was taken. Specifically, for each *All of Us* participant an outer time boundary was taken from the collection of all drug exposures that were statin drug exposures. If this window of time overlapped the day on which a particular lipids measurement occurred, then that measurement was adjusted for status use. We did not correct for missingness of drug exposure data in this study, but it is a known issue worth addressing in future similar studies. For full details, please see [01_aou_lipids_phenotype.ipynb](https://github.com/all-of-us/ukb-cross-analysis-demo-project/blob/main/aou_workbench_siloed_analyses/01_aou_lipids_phenotype.ipynb).

#### **Adjustment of lipid levels for statin use**

For both UK Biobank and *All of Us*, TC and LDL-C measurements were adjusted for lipid lowering medications as previously done^5^ (**Supplemental Fig. 7**).

1. LDL-C adjustment based on TG/LDL-C values
   - If TG > 400, then LDL-C = NA
   - If TG is null, then LDL-C = LDL-C
   - If LDL-C < 10, then LDL-C = NA
2. LDL-C and TC adjustment based on Statin (Lipid lowering medication)
   - If STATIN is used, LDL-C_ADJ = LDL-C/0.7
   - If STATIN is used, TOTAL_ADJ = TC/0.8
3. TG adjustment
   - TG_LOG = log(TG)

***Supplemental Fig. 7****: Formula used for lipid level adjustment per statin use.*

### **Lipids GWAS analysis method**

For both UK Biobank and *All of Us*, the lipid phenotypes were inverse rank normalized the residuals, scaled by the standard deviation and adjusted for the covariates. We included PC1-10, age, age^2^ and sex at birth as covariates in our study ^6^. For the pooled analysis an additional covariate of ‘cohort’ was added to mitigate batch effects. The PCs were generated using a high-quality LD pruned set of variants using Plink ^7^.

Single variant genome wide association studies (GWAS) were carried out using REGENIE v2.2.4 ^8^. For the pooled analysis we used 299,265 samples for which lipid phenotype data was available for our analysis. We implemented REGENIE Step1 NULL model generation using 313,211 quality-controlled variants with a minor allele count (MAC) of 100. We applied the leave one chromosome out (LOCO) method for GWAS while adjusting for the covariates stated above. We used variant and sample missingness at 10% followed by Hardy-Weinberg equilibrium p-value not exceeding 1x10^-15^ for the genome wide associations. In total we tested the association of 2,135,845 variants with four lipids. For UK Biobank analysis we used 190,982 samples for which lipid phenotype data was available for our analysis. We implemented REGENIE Step1 NULL model generation using 267,898 quality-controlled variants with an MAC of 100. We applied the LOCO method for GWAS while adjusting for the covariates stated above. We used variant and sample missingness at 10% followed by Hardy-Weinberg equilibrium p-value not exceeding 1x10^-15^ for the genome wide associations. In total we tested the association of 2,037,169 variants with four lipids. For *All of Us* analysis we used 37,754 samples for which lipid phenotype data was available for our analysis. We implemented REGENIE Step1 NULL model generation using 179,349 quality-controlled variants with an MAC of 100. We applied the LOCO method for GWAS while adjusting for the covariates stated above. We used variant and sample missingness at 10% followed by Hardy-Weinberg equilibrium p-value not exceeding 1x10^-15^ for the genome wide associations. In total we tested the association of 789,179 variants with four lipids.

We carried out meta-analysis of the GWAS results from both the cohorts using the METAL package ^9^. We used the Standard Error scheme, where the methods weights effect size estimates using the inverse of the corresponding standard errors. We used the standard error (SE) method to account for the differences in allele frequencies between studies, only using sample size for the weight can be suboptimal in terms of power of the meta-analysis. **Supplemental Table 1** provides a summary of lipid GWAS results. **Supplemental Tables 2 & 3** and **Supplemental Figure 8** provide a summary of significant variants from lipid GWAS. **Supplemental Figure 9** compares the LOG10P values for variants examined in both approaches, but meeting the genome-wide significance level (p<5E-08) in only one approach.

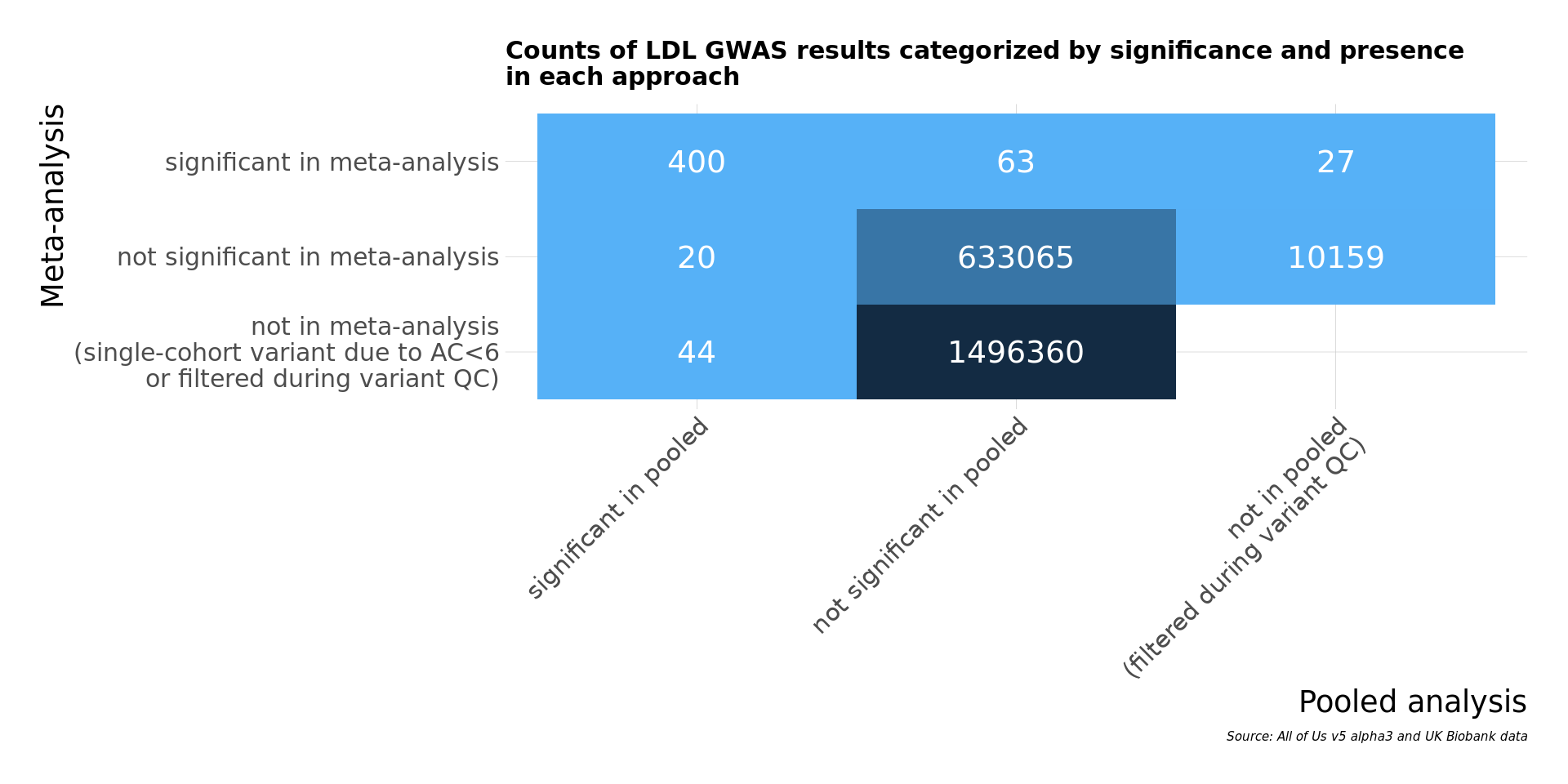

***Supplemental Fig. 8****: LDL-C variant counts categorized by their significance, and in which result sets they occurred. The 27 significant results from the meta-analysis not present in pooled were all removed during the variant QC process via the Hardy Weinberg equilibrium filter.*

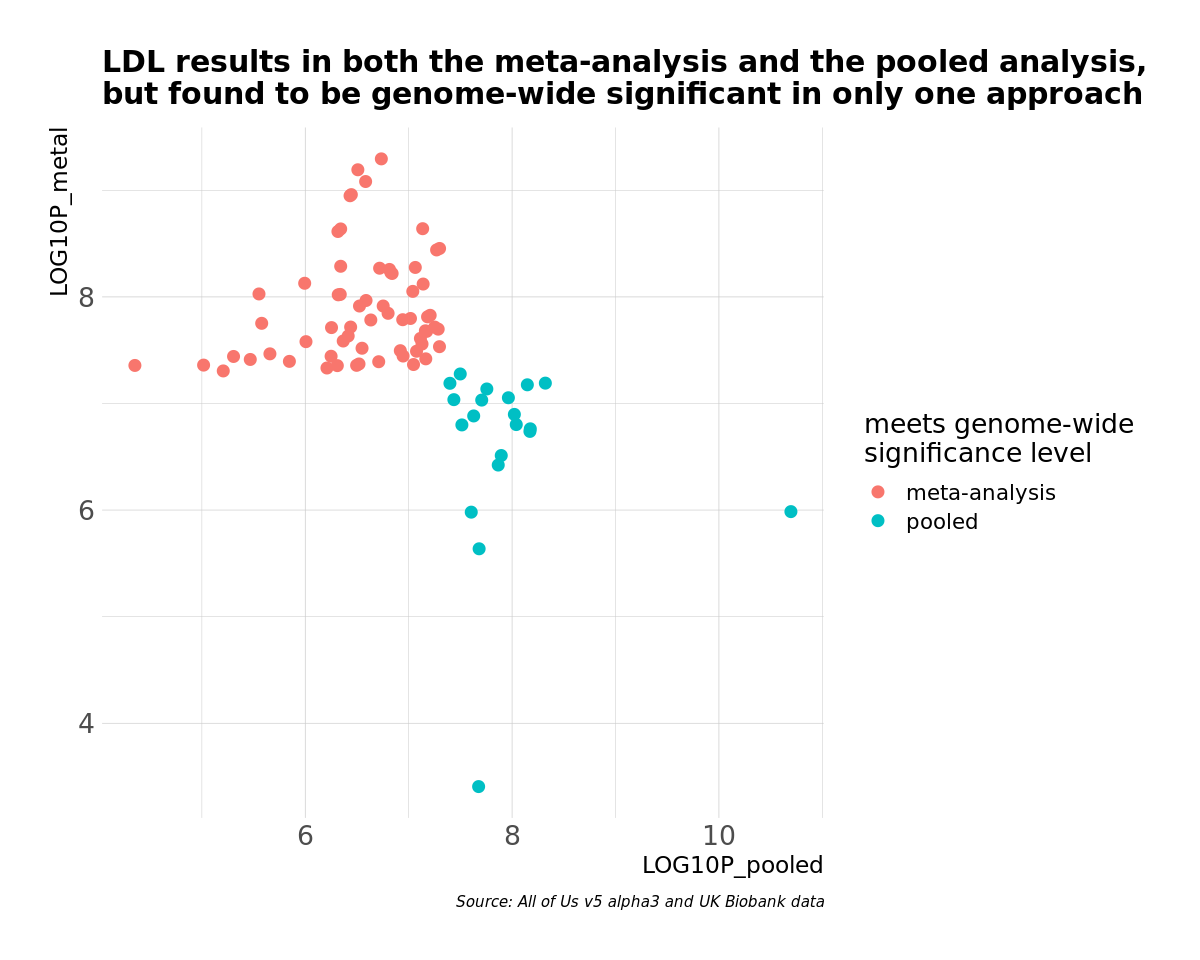

***Supplemental Fig. 9****: LOG10 (P-values) for LDL-C GWAS found to meet the genome-wide significance level in one approach, but not the other. Most LOG10 (P-values) for the method not meeting the genome-wide significance are close to that hard cut off of p<5E-08.*

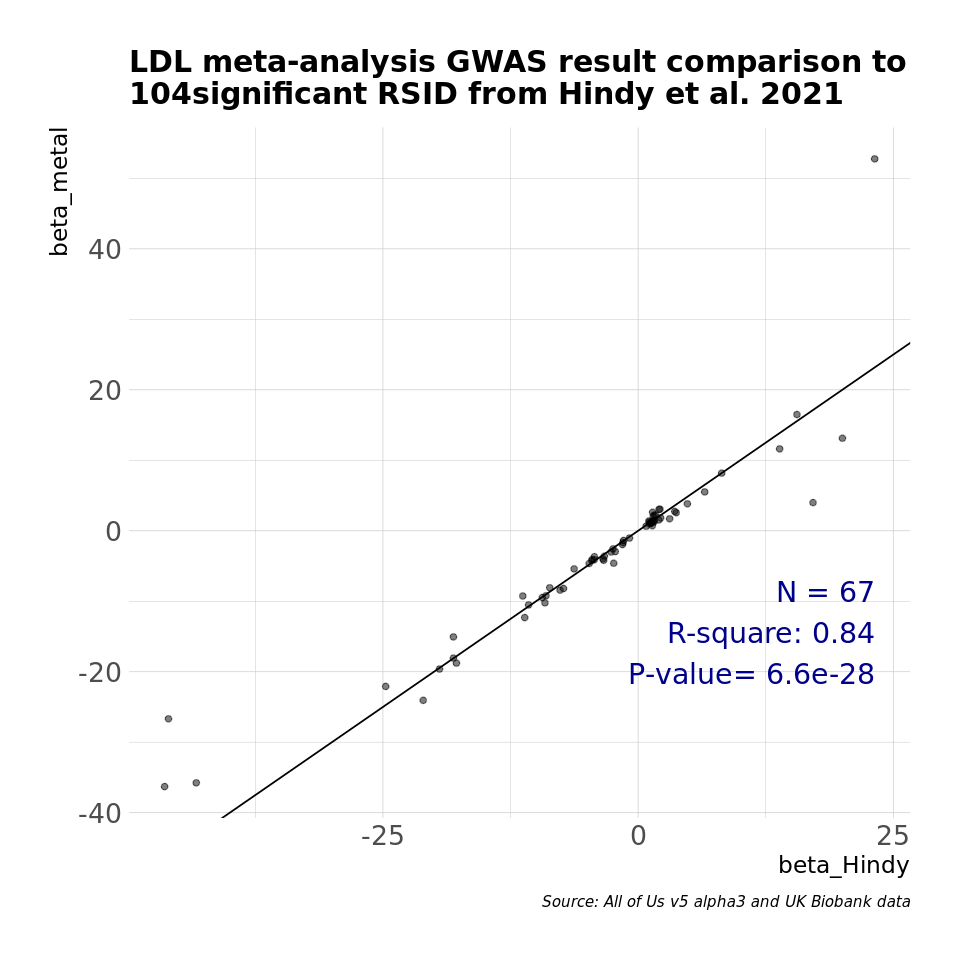

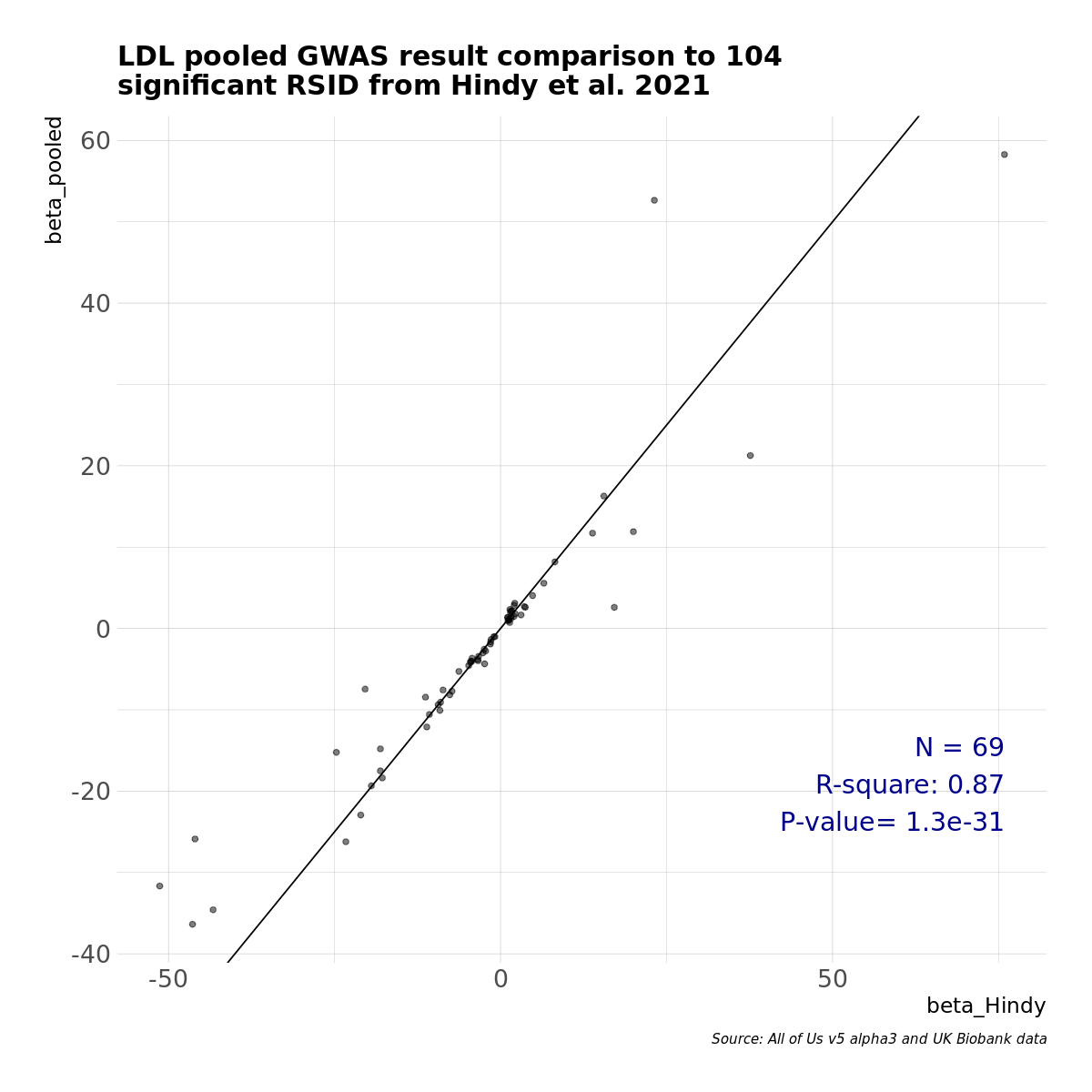

***Supplemental Fig. 10a****: Comparison of results against published whole exome dataset*^10^*. Left: meta-analysis results Right: pooled results*

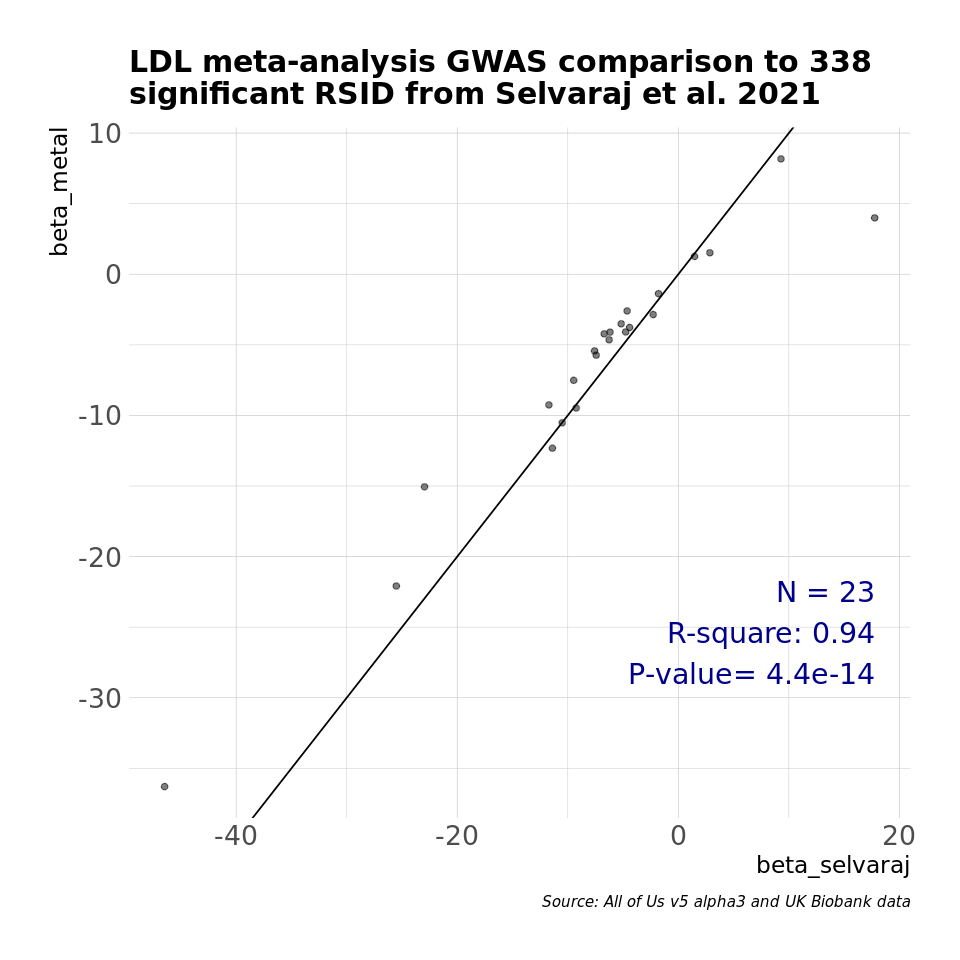

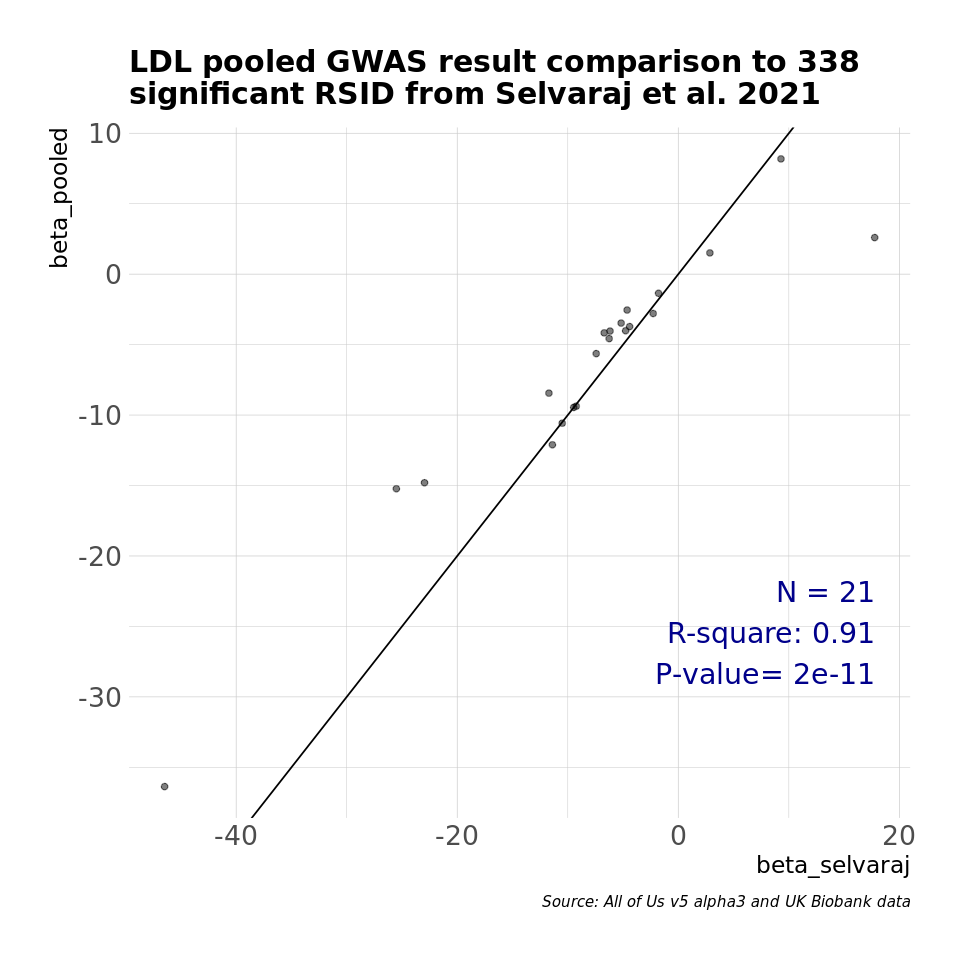

***Supplemental Fig. 10b****: Comparison of results against published whole genome dataset* ^11^*. Left: meta-analysis results Right: pooled results*

In **Supplemental Fig. 11** note that the duplicative computational steps performed in the meta-analysis yield a total of 32 arrows, where the arrows are the inputs/outputs to each computational step. For the pooled approach, we have a total of 19 arrows, which is still a fair bit of complexity, but all computational steps are unique.

If a future analysis utilized data from three distinct TREs, the meta-analysis would yield 32 + 16 = 48 arrows and the pooled approach would yield 20 + 4 = 24 arrows. For N distinct TREs, the meta-analysis would yield 16*N arrows and the pooled approach would yield 12 + 4*N arrows.
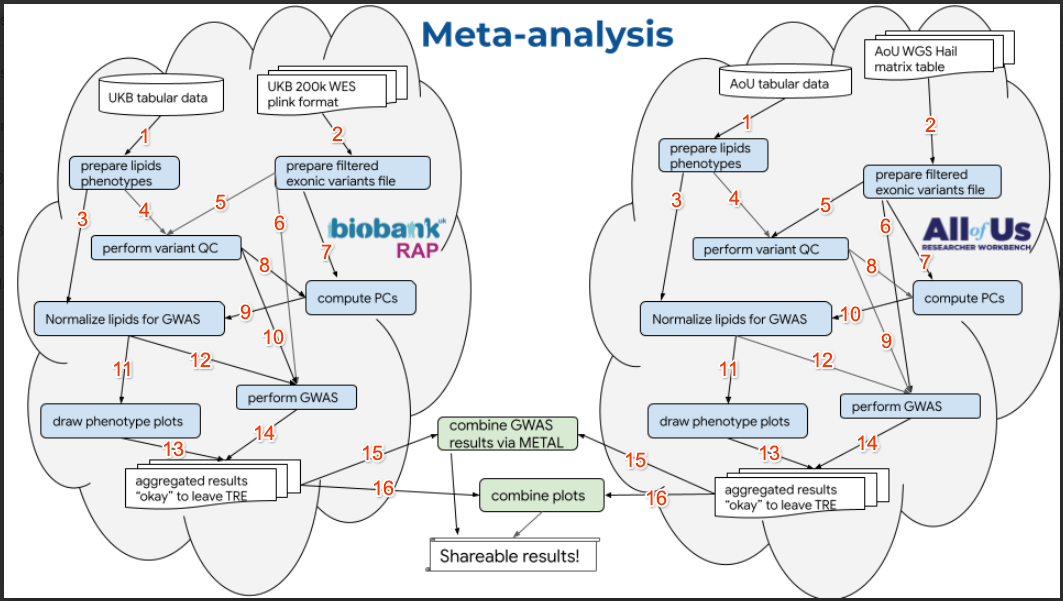

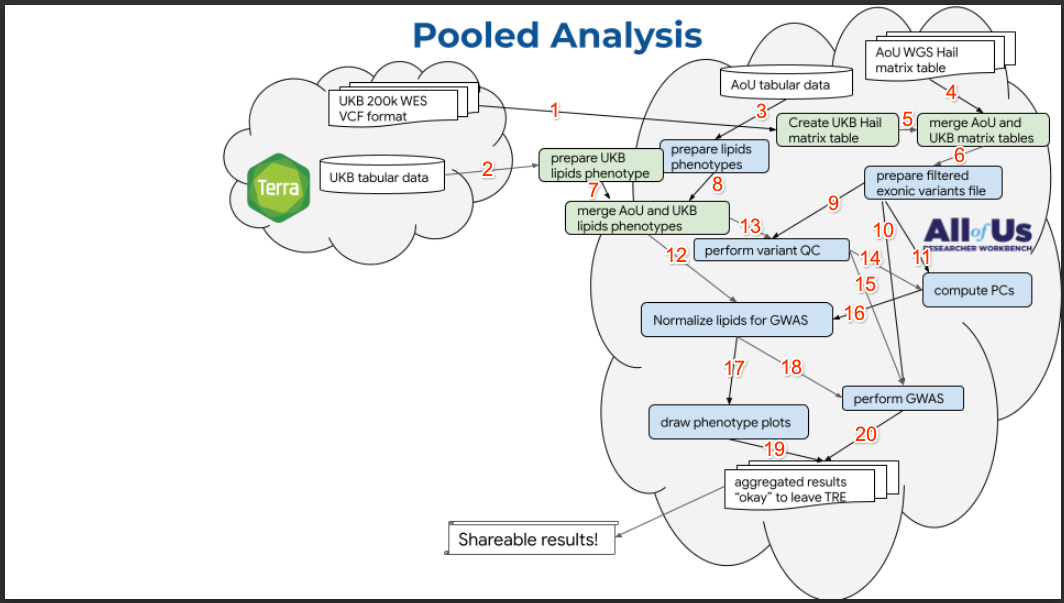

***Supplemental Fig. 11****: Steps involved in the meta- and pooled analyses.*

### **Ancestry and Functional Analysis**

We curated variant information from the [gnomAD database](https://gnomad.broadinstitute.org/) to understand the enrichment of functional categories and ancestral distribution for the significant variants identified from our analysis. We used the population maximum allele frequency (popmax) as defined by gnomAD ^12^, where ancestral groups were defined as African/African American, East Asian, European, Latino/Admixed American, and South Asian (**Supplemental Tables 2, 3, and 4**). We used scaled Combined Annotation Dependent Depletion (Phred-like CADD) scores ^13^ and annotations from variant effect predictor (VEP) ^14^ to define functional importance of the variants.

**References**

1.	Szustakowski, J. D. *et al.* Advancing human genetics research and drug discovery through exome sequencing of the UK Biobank. *bioRxiv* (2020) doi:10.1101/2020.11.02.20222232.

2.	Yun, T. *et al.* Accurate, scalable cohort variant calls using DeepVariant and GLnexus. *Bioinformatics* (2021) doi:10.1093/bioinformatics/btaa1081.

3.	Allen NE, et al. Approaches to minimising the epidemiological impact of sources of systematic and random variation that may affect biochemistry assay data in UK Biobank. Wellcome Open Res. 2021 Jan 4;5:222. doi: 10.12688/wellcomeopenres.16171.2.

4.	Rugge, B. *et al.* *Lipid Conversion Factors*. (Agency for Healthcare Research and Quality (US), 2011).

5.	Data Methods – All of Us Research Hub. https://www.researchallofus.org/data-tools/methods/.

6.	Natarajan, P. *et al.* Deep-coverage whole genome sequences and blood lipids among 16,324 individuals. *Nat. Commun.* **9**, 3391 (2018).

7.	Purcell, S. *et al.* PLINK: a tool set for whole-genome association and population-based linkage analyses. *Am. J. Hum. Genet.* **81**, 559–575 (2007).

8.	Mbatchou, J. *et al.* Computationally efficient whole-genome regression for quantitative and binary traits. *Nat. Genet.* **53**, 1097–1103 (2021).

9.	Willer, C. J., Li, Y. & Abecasis, G. R. METAL: fast and efficient meta-analysis of genomewide association scans. *Bioinformatics* **26**, 2190–2191 (2010).

10.	Hindy, G. *et al.* Rare coding variants in 35 genes associate with circulating lipid levels-A multi-ancestry analysis of 170,000 exomes. *Am. J. Hum. Genet.* **109**, 81–96 (2022).

11.	Selvaraj, M. S. *et al.* Whole genome sequence analysis of blood lipid levels in >66,000 individuals. *bioRxiv* 2021.10.11.463514 (2021) doi:10.1101/2021.10.11.463514.

12.	Karczewski, K. J. *et al.* The mutational constraint spectrum quantified from variation in 141,456 humans. *Nature* **581**, 434–443 (2020).

13.	Rentzsch, P., Witten, D., Cooper, G. M., Shendure, J. & Kircher, M. CADD: predicting the deleteriousness of variants throughout the human genome. *Nucleic Acids Res.* **47**, D886–D894 (2019).

14.	McLaren, W. *et al.* The Ensembl Variant Effect Predictor. *Genome Biol.* **17**, 122 (2016).
